# Human circulatory system response to changes in alveolar pressure and lung volume

**DOI:** 10.1101/2024.03.16.585354

**Authors:** Yu.S. Semenov, A.I. Dyachenko, I.S. Melnikov, R.N. Zaripov

## Abstract

The response of hemodynamic parameters in healthy volunteers to changes in alveolar pressure and lung volume was studied by noninvasive methods during respiratory maneuvers similar to Valsalva and Müller maneuvers (in a sitting position and lying on the back horizontally). The following lung volumes and values of pressure (relative to atmospheric pressure) were considered in various combinations: total lung capacity, functional residual capacity, residual volume, -30, -15, 0, +15, +30 mmHg. Changes in hemodynamic parameters averaged over the duration of a maneuver were studied (the duration of a maneuver was 30 s). Changes in alveolar and, accordingly, intrathoracic pressure influenced hemodynamic more strongly than changes in lung volume or body position. Stroke volume decreased with increasing alveolar pressure and increased with decreasing pressure regardless of lung volume and body position; changes ranged from -35 to +15 ml. The effect of changes in alveolar pressure was more pronounced in a sitting position. Heart rate increased with increasing alveolar pressure (up to +20 bpm) but changed little with decrease in pressure. Mean arterial pressure decreased with decreasing alveolar pressure regardless of lung volume and body position; with increasing alveolar pressure, the result depended on lung volume. When performing maneuvers at total lung capacity, mean arterial pressure remained below baseline values, in other cases it increased. Changes in mean arterial pressure were within ±20 mmHg. Regardless of lung volume and body position, total peripheral resistance decreased with decreasing alveolar pressure and increased with increasing alveolar pressure; the range of changes in total peripheral resistance was -0.3 to +0.7 mmHg·s/ml.

## Introduction

It is well known that breathing affects the systemic circulation. One of the main factors altering blood circulation is intrathoracic pressure. Intrathoracic pressure changes during the respiratory cycle, during various breathing maneuvers, and also in certain lung diseases.

The effect of breathing on blood circulation, caused by fluctuations in intrathoracic and alveolar pressure, makes it possible to use breathing maneuvers to diagnose various human diseases. Some of the most famous and widespread are two breathing maneuvers-antipodes: the Valsalva test (an increase in pressure in airways for 15–20 seconds created by the effort of expiratory muscles of the tested person) and the Müller test (a decrease in pressure created by inspiratory muscles).

Since there is no airflow in the airways when performing Valsalva or Müller maneuvers or similar ones, oral and alveolar pressures are equal, which makes the mentioned maneuvers a convenient tool to study the interaction of breathing and circulation. Intrathoracic and alveolar pressures when performing Valsalva or Müller maneuvers differ by the value of the lung elastic recoil, which depends on lung volume.

The Valsalva maneuver is used to diagnose many human diseases, using the peculiarities of the circulatory reaction in the known four phases of the reaction [Nishimura and Tajik, 1986; Pstras et al., 2016]. The dynamics of the circulatory response to the Müller maneuver is described, for example, in [Hemalatha and Manivannan, 2011].

Intrathoracic pressure deviations have clinical importance. They are greatly increased when the airway resistance increases, for example, during bronchial asthma attacks or during obstructive apnea [Pressman et al., 2016]. Obstructive sleep apnea is also associated with the frequent occurrence of atrial fibrillation and other disorders of the cardiovascular system (CVS) [Friend et al., 2022; Linz et al., 2021; Pressman et al., 2016]. Large negative pressure creates a background for pulmonary edema [Lemyze and Mallat, 2014], and large positive pressure limits venous blood return to the heart, thereby reducing stroke volume [Nishimura and Tajik, 1986].

It should be noted that not only breathing affects blood circulation, but also blood circulation affects breathing. For example, the influence of the function and dysfunction of heart ventricles on pulmonary circulation and gas exchange in the lungs is the object of study [Comunale et al., 2021; Luo et al., 2011].

To assess how various disorders of respiratory mechanics can affect circulation, it is important to understand what changes occur in the CVS under different combinations of lung volumes and alveolar pressure. Thus, the question of the nature of processes in the CVS in response to changes in lung air pressure has not only theoretical but also practical importance.

The quantitative description of the dynamics of CVS parameters during Valsalva and Müller maneuvers cannot be called complete and systematic, although repeated attempts were made to use the quantitative description for diagnostic purposes. For example, there are values of some parameters in four phases of the Valsalva test [Greenfield et al., 1984]. And in [Lu et al., 2001], available notions about CVS regulation were sufficient to reproduce in a mathematical model the normal response of arterial pressure, heart rate (HR), and cardiac output (CO) to the Valsalva maneuver with different increases in oral pressure.

Considerable attention was paid to the effect of maneuvers on the distribution of blood filling and blood flow in the lungs [Bake, 1971]. Alveolar pressures of +20 and +40 mmHg significantly increased the irregularity, and pressures of -5 to -40 mmHg significantly decreased the irregularity of blood flow. Bake [Bake, 1971] associated these results with changes in venous return. Studies of dynamics of the lung diffusion capacity during maneuvers allowed the authors to estimate the dynamics of blood volume in pulmonary capillaries [Smith and Rankin, 1969]: the volume increased by 20% during Müller maneuver, and it decreased at 15th second of Valsalva maneuver by approximately 1.5 times.

Although some hemodynamic effects of changes in intrathoracic pressure and lung volume are known, there is no clear separation of contributions of intrathoracic pressure and lung volume to changes in hemodynamics. Separation of the contributions is important because corresponding effects are determined by vessels of pulmonary circulation, but by its different parts: alveolar and extra-alveolar vessels. Alveolar pressure acts on the external surface of alveolar vessels of the lungs, so primarily this pressure determines the condition of alveolar vessels. Tissue pressure strongly depends on lung volume and affects the external surface of extra-alveolar vessels. Therefore, one can expect that resistance and blood volume in extra-alveolar vessels also strongly depend on lung volume.

As a result, two parameters (current values of alveolar pressure and lung volume) set the external influence on pulmonary vessels. However, these two parameters describe only external influence on pulmonary vessels. Another important parameter influencing pulmonary hemodynamics is the blood filling of vessels related to blood pressure inside them. In experiment, pressure inside pulmonary vessels and their blood filling can be changed noninvasively. To do this, it is sufficient to change the position of the body relative to the horizon.

**The aim of our work** was to obtain quantitative information about the response of human system hemodynamics to changes in alveolar pressure and lung volume in different body positions.

The results of this study will help better understanding of mechanisms underlying the interaction between breathing and circulation, will provide information for the development of diagnostic techniques and methods, and will be useful for those who are engaged in modeling the cardiorespiratory system.

## Materials and Methods

We used a reduction and an increase in mouth and alveolar pressures at fixed lung volume to separate the influence of alveolar pressure and lung volume on circulation. We compared CVS parameters at identical lung volumes but different mouth pressures, as well as at identical mouth pressures but different lung volumes. At a specified lung volume, volunteers attempted to inhale or exhale through a closed mouthpiece while attempting to maintain a prescribed level of mouth pressure for 30 seconds. These breathing maneuvers, which are close to the Valsalva and Müller maneuvers but better standardized, were performed in supine horizontal position and in a sitting position. Changes in body position lead to changes in baseline level (i.e. before the maneuver) of arterial pressure in carotid and aortic reflexogenic zones, intrathoracic pressure, lung blood filling and lung volumes, characteristics of external influence on pulmonary vessels, which can markedly change the CVS reactions.

Due to time constraints, the study was divided into three series performed with different, but similar in basic characteristics (age, height, weight) groups of volunteers; some of the volunteers participated in two or three series. All series followed the same scheme. Only the values of alveolar pressures, lung volumes, and body positions differed. In order to assess the reproducibility of reactions between the three groups of volunteers, some of the maneuver variants were performed by all three groups of volunteers.

Each volunteer performed several dozens of breathing maneuvers, each lasting 30 seconds. Two minutes between two consecutive maneuvers were allotted for rest and recovery; recovery was monitored by the return of physiological parameters to baseline (initial) values. A nose clip was used during a maneuver, which was removed during the rest period. Volunteers performed maneuvers at different values of lung volume, two body positions (sitting or lying on the back), and different airway pressures. To exclude the influence of possible fatigue and prolonged physiological reactions caused by breath-holding during maneuvers, the sequence of lung volume and mouth pressure combinations for each volunteer was chosen randomly. For consecutive maneuvers, the same combination was not repeated unless such repetition was a consequence of random sequence. A pseudorandom number generator formed initial sequence, then corrected so that each combination of lung volume and mouth pressure occurred the same number of times in the sequence. The entire planned set of maneuvers for each of the body positions was performed in a single block without changing the body position. Volunteer’s instruction prohibited any preparatory actions (hyperventilation, single deep breaths, etc.) before performing the maneuver, and the execution of the instruction was strictly controlled. If for any reason the maneuver was not successful (the volunteer failed to keep the pressure in the airways for 30 seconds or made a mistake with its value, malfunction of the recording equipment, etc.), this variant of the maneuver was repeated later, randomly including the repetition in the planned sequence. For any of the maneuver variants, at least 4 “clean” repetitions (total in a series, not in a row) were collected for each volunteer, by which the average individual reaction to the given maneuver variant was calculated in the subsequent analysis of measurement results. After the completion of planned set of maneuvers, volunteers rated its difficulty on a scale from 0 to 7 (where 0 was “no effort”, 1 was “very easy”, 2 – “somewhat easy”, 3 – “moderate”, 4 – “somewhat hard”, 5 – “hard”, 6 – “very hard”, 7 – “very, very hard, no endurance for long”) and marked and rated (the same scale) the most difficult variant of the maneuvers.

Each volunteer was trained to perform maneuvers before the very beginning of the planned set of maneuvers.

Volunteers performed maneuvers at the depth of full deep exhalation (residual volume, RV), at the level of functional residual lung capacity (FRC), and at the height of full deep inhalation (total lung capacity, TLC). The volunteers’ task was to hold the specified pressure in closed mouthpiece as accurately as possible for 30 seconds while attempting to inhale (RV, FRC maneuvers) or exhale (TLC, FRC maneuvers) through the mouthpiece. We used the following pressure levels (relative to atmospheric pressure): +30 mmHg, +15 mmHg, -15 mmHg, -30 mmHg, 0 (atmospheric pressure in the airways, epiglottis open). Hereinafter pressure level and lung volume indicate the type of maneuver, e.g. +30_TLC – maneuver performed at total lung capacity at pressure in the mouthpiece of +30 mmHg. The 0_FRC maneuver was used as a control to assess the contribution of changes in blood gas tensions due to 30-second apnea to the observed cardiovascular response. Information on the set of conditions for the maneuvers is given in Table 1 (11 variants of maneuvers for each of the body position).

**Table 1.** Variants of combinations of lung volume, alveolar pressure (relative to atmospheric pressure), and the body position used during breathing maneuvers for three groups of volunteers (explanations in the text). RV – residual volume, FRC – functional residual capacity, TLC – total lung capacity; I, II and III – numbers of volunteer groups (study series), shaded cells – “forbidden” combinations, which cannot be realized when performing voluntary breath-holds.

| Lying position |  | Alveolar pressure, mmHg |  |  |  |  |
| --- | --- | --- | --- | --- | --- | --- |
|  |  | +30 | +15 | 0 | -15 | -30 |
| Lung volume | RV |  |  | II | I | I, II, III |
|  | FRC | I | I | III | I | I |
|  | TLC | I, II, III | I | II |  |  |
| Sitting position |  | Alveolar pressure, mmHg |  |  |  |  |
|  |  | +30 | +15 | 0 | -15 | -30 |
| Lung volume | RV |  |  | III | I | I, III |
|  | FRC | I | I | III | I | I |
|  | TLC | I, III | I | III |  |  |

Before and during maneuvers, as well as after them (during rest periods between maneuvers), main CVS parameters were continuously measured: heart rate (HR), systolic (SAP), diastolic (DAP) and mean (MAP) blood pressure (for each cardiac cycle using a modified Peňáz method) [Wesseling, 1984], stroke volume (SV, for each cardiac cycle), cardiac output (CO), total peripheral resistance (TPR), periods of isovolumetric contraction and ventricular ejection. Stroke volume was estimated according to the parameters of pulse pressure wave registered by the Peňáz method and the known population-average mechanical characteristics of the vascular bed [Jansen et al., 2001; Wesseling et al., 1993] (Finometer Pro, Finapres Medical Systems BV, the Netherlands). Finometer Pro software also evaluated baroreflex sensitivity (BRS). BRS was measured as the slope of the dependence of the R-R interval duration versus mean arterial pressure [La Rovere et al., 2008]. Periods of isovolumetric contraction and ventricular ejection were measured by thoracic rheography [Kubicek et al., 1970]. For recovery evaluation after the maneuver completion, oxygen and carbon dioxide tensions in the skin (left subclavicular region) and arterial blood oxygen saturation (pulse oximetry method, SpO_2_) were continuously measured using a transcutaneous monitor (TCM4, Radiometer Medical AsP, Denmark). Due to a notable response delay of the transcutaneous method, its results were not analyzed in detail and were used only during measurements to assess recovery. The next maneuver was performed only after the initial values of the parameters were restored. If the recovery occurred earlier than the planned two minutes, the participants waited for the completion of the 2-minute break before performing the next maneuver.

In study series II and III, we also used bioimpedance technique to assess blood filling of the right lung (measurements were performed by tetrapolar scheme). Electrodes were placed along the right midclavicular and scapular lines. The upper current electrode was placed on the upper surface of the shoulder, the upper potential electrode was placed on the front torso surface 10 cm below the upper current one, the lower potential electrode was placed on the back at the level of the lower edge of the lungs, and the lower current electrode was located 10 cm below the lower potential one [Orkvasov et al., 2011]. The lower edge of the right lung was determined by percussion. Since the bioimpedance measurement area includes the right lung, impedance depends not only on the amount of fluid in the measured volume but also on the current lung volume (amount of gas in lungs), as well as on the geometry of the chest (the distance between the electrodes changes together with it). To asses lungs hydration and its changes during maneuvers performing, we used R5 to R50 ratio (R5 and R50 are the real parts of impedance at 5 kHz and 50 kHz, respectively). The ratio does not depend on the geometry of the chest: as the same electrodes are used for measurements, the distance between them does not affect R5/R50 value. Since gas in lungs is an insulator, R5/R50 is mainly affected by the amount of fluid in the measured volume (more accurate, by the ratio of amounts of intra- and extracellular fluid). When performing breathing maneuvers, only the amount of extracellular fluid, mainly blood, can change. In [Orkvasov et al., 2011], it is shown that the ratio R5/R50 (lung hydration index) decreases when the amount of fluid in lungs increases, and vice versa. In healthy men R5/R50 ranges from 1.19 to 1.27 (when measured in supine position).

Rheographic measurements were performed at 40 kHz (RPKA2-01, STC «Medass», Russia), and bioimpedance measurements were performed at 5 and 50 kHz (Sprut-2m, STC «Medass», Russia).

The volunteer controlled pressure in the mouthpiece and in the airways by adjusting the force developed by respiratory musculature. The current pressure was displayed on a display in the volunteer’s field of view. The mouthpiece had a leakage to avoid a situation in which the volunteer closed the epiglottis and created pressure in the mouthpiece solely by contraction of the facial, laryngeal, and mandibular muscles (with alveolar pressure not equal to mouth pressure). The created leakage was sufficient for the rapid depletion of air in the oral cavity and not enough to noticeably (no more than 5– 10% of the current value) change the lung volume during the maneuver. To control the position of the epiglottis when volunteer was performing maneuvers at atmospheric pressure in the airways, we used pulse oscillometry method (Jaeger MasterScreen IOS, VIASYS Healthcare GmbH, Germany). Pulse oscillometry allowed us to measure airway resistance, which increases manifold even when the epiglottis is partially closed.

Lung volumes (RV, TLC, FRC) and maximal voluntary efforts of respiratory muscles during inspiration (Pins) and expiration (Pexp) were measured (Jaeger MasterScreen PFT, VIASYS Healthcare GmbH, Germany) in volunteers before starting the study.

Nine healthy men of average build aged 20–36 years (mean age ± SD = 26±5) participated as volunteers in study series I. Their TLC, RV, FRC, Pins and Pexp (mean ± SD) were 7790.0±984.3 ml, 1584.4±335.9 ml, 4593.3±938.1 ml, 12.3±1.9 kPa and 19.0±2.6 kPa respectively, data obtained in a sitting position. Each volunteer performed 64 maneuvers (32 in a supine position, 32 in a sitting position). Each of the 8 maneuver variants (combinations of the body position, lung volume, and alveolar pressure) was successfully repeated 4 times. A total of 9x64=576 maneuvers were analyzed.

Ten healthy men of average build aged 20–35 years (mean ± SD = 26±5) volunteered for study series II. Their TLC, RV and FRC (mean ± SD) were 7708.0±512.6 ml, 1668.0±185.1 ml and 4247.0±559.9 ml respectively, data obtained in a sitting position. Each volunteer performed 20 maneuvers (all in a supine position). Each of the 4 maneuvers (combinations of lung volume and alveolar pressure) was successfully repeated 5 times. A total of 10x20=200 maneuvers were analyzed.

Nine healthy men of average build aged 21–31 years (mean age ± SD = 27±4) participated in study series III. Their TLC, RV, FRC, Pins and Pexp (mean ± SD) were 7751.1±649.8 ml, 1567.8±236.5 ml, 4572.2±547.0 ml, 10.0±1.7 kPa and 11.4±3.1 kPa respectively, data obtained in a sitting position. Each volunteer performed a minimum of 40 maneuvers (18 in a supine position, 22 in a sitting position). Each of the 8 maneuver variants (combinations of body position, lung volume, and alveolar pressure) was successfully repeated 6 times (except for the maneuver variants used to compare groups: -30_RV sitting and +30_TLC sitting, wherein there were 2 repetitions). A total of 9x40=360 maneuvers were analyzed.

A total of 1136 maneuvers with 22 different variations were analyzed for the entire study.

In accordance with the Declaration of Helsinki, written informed consent was obtained from all participants and the studies were approved by the Bioethical Commission of the Institute of Biomedical Problems of the Russian Academy of Sciences (protocol No. 395 of May 27, 2015).

Data analysis for all study series was performed in the same way. All obtained time series of data with values of synchronously registered physiological parameters were split into fragments (frames) corresponding to the performance of an individual maneuver. The beginning and the end of the maneuver were marked by the signal representing pressure in the mouthpiece cavity.

For the maneuvers performed at atmospheric pressure in the respiratory tract, the mark of the beginning and the end of the maneuver was also entered in the channel of the pressure signal and was performed by applying a constant voltage (outside the range of pressure signal changes) throughout the maneuver. Researchers manually set the mark according to the spirometer data used in the pulse oscillometry method. In this method of marking, an error of 0.5–1 s was possible, but this synchronization accuracy was sufficient for subsequent analysis.

After splitting the recordings and sorting the frames by combinations of lung volume, alveolar pressure, and body position, we calculated average curves reflecting changes in physiological parameters for each maneuver variant and each volunteer, averaging was performed by repetitions of the maneuver. The resulting curves included 30 seconds before the maneuver, the section corresponding to the execution of the maneuver, and two minutes after the completion of the maneuver, total frame duration was 3 minutes. Before averaging, the baseline value was subtracted from the parameter values. Average parameter value calculated from the 20-second segment of the recording before the beginning of the maneuver was considered as the baseline. In this way, individual average curves representing changes of physiological parameters from their baseline during certain variant of the maneuvers were found. Then, for each of the maneuver variants, these curves were averaged over the group of volunteers.

After the calculation of group average curves, we determined the average value of deviation over 26 seconds of the maneuver for each of the parameters (2 seconds at the beginning and 2 seconds at the end of the maneuver were discarded). These average deviation values were the final result of data processing.

Deviations that differed from zero by more than 2SD were considered statistically significant.

## Results

All volunteers successfully performed the planned maneuvers. On a scale from 0 to 7 volunteers rated the difficulty of proposed breathing maneuvers as 2–5. The most frequent answer was 4 (“somewhat hard”). However, it was noted that some variants of maneuvers were very difficult, they were evaluated at 5–7 points, with different volunteers marked as very difficult different variants of maneuvers.

Arterial blood oxygen saturation (SpO_2_) was virtually unchanged during any of the maneuvers. An apnea-induced decrease began only a few seconds before the end of the maneuver and was observed only for RV maneuvers and FRC maneuver variants with a decrease in alveolar pressure. The maximum decrease was not over 3–5% and was observed about 10 seconds after the end of the maneuver. During the performance of any maneuver variant, SpO_2_ even slightly increased (by 0.1–1%). With any of maneuver variant, transcutaneous CO_2_ tension changed no more than several mmHg, the response (decrease) developed after the end of the maneuver and was much more pronounced in maneuvers with positive alveolar pressure.

For all recorded physiological parameters and practically all maneuver variants, there was a significant (exceeding 2 SD interval) change of parameter values from baseline.

Individual differences in the response of CVS to breathing maneuvers exceeded the differences between attempts to perform the same variant of the maneuver by the same volunteer. All volunteers had unidirectional changes in physiological parameters when performing the same variant of the maneuver.

A two-minute break between consecutive maneuvers was sufficient in all cases to recover from the previous maneuver.

For most of the parameters during maneuvers with any combination of lung volume, alveolar pressure and body position, we observed complex multiphase dynamics of parameter changes. For example, for mean arterial pressure (MAP) and heart rate (HR), the pattern of response to positive alveolar pressures was similar to that observed by many researchers studying the response of CVS to the Valsalva maneuver (Fig. 1). When performing the +30_TLC maneuver, which is similar in characteristics to the Valsalva maneuver, four phases of CVS response described in the literature [Lu et al., 2001; Pstras et al., 2016] could be distinguished on the curves.

**Fig. 1.**
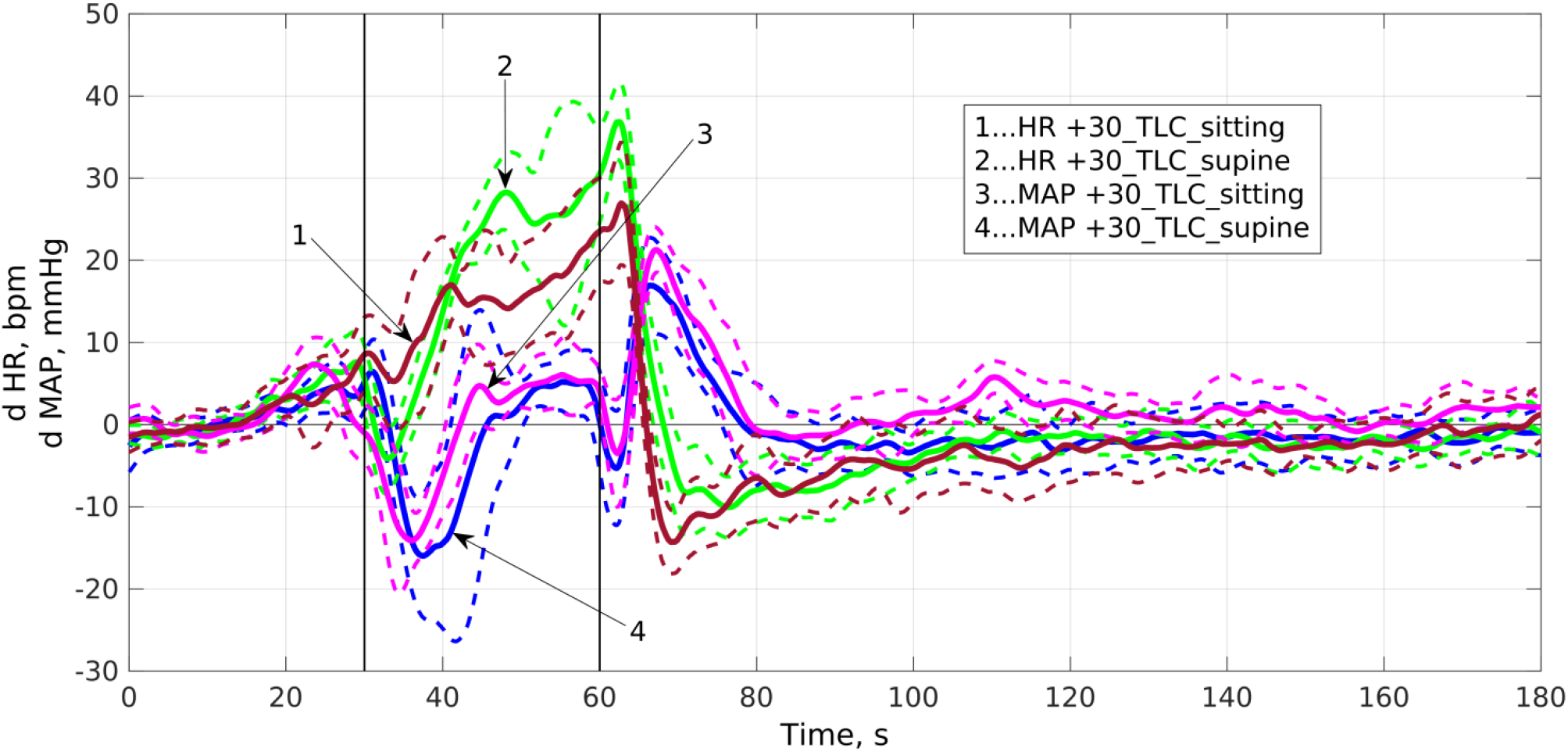
Example of the dynamics of heart rate (HR) and mean arterial pressure (MAP) during the +30_TLC maneuver. Group I average curves are shown. Vertical lines indicate the start and the end points of the maneuver. Dashed lines correspond to the ±SD range. Prefix “d” denotes parameter changes from its baseline value.

The multiphase response is associated with the involvement of various physical processes and the activation of various physiological circulatory regulation circuits, each of which has its own amplitude and time characteristics and can act in different ways on the observed parameter. The multiphase response complicates the interpretation of obtained results. Parameters that are more convenient for analysis are characterized by a trapezoid response (Fig. 2). Such parameters with simple dynamics of deviation from the baseline include stroke volume and ventricular ejection period. The trapezoidal shape did not change during performance of the maneuver with any combination of lung volume, alveolar pressure or body position. For uniformity, despite the possibility of distinguishing several phases in some cases, all measured physiological parameters were processed in the same way by averaging deviation from the baseline over duration of the maneuver.

**Fig. 2.**
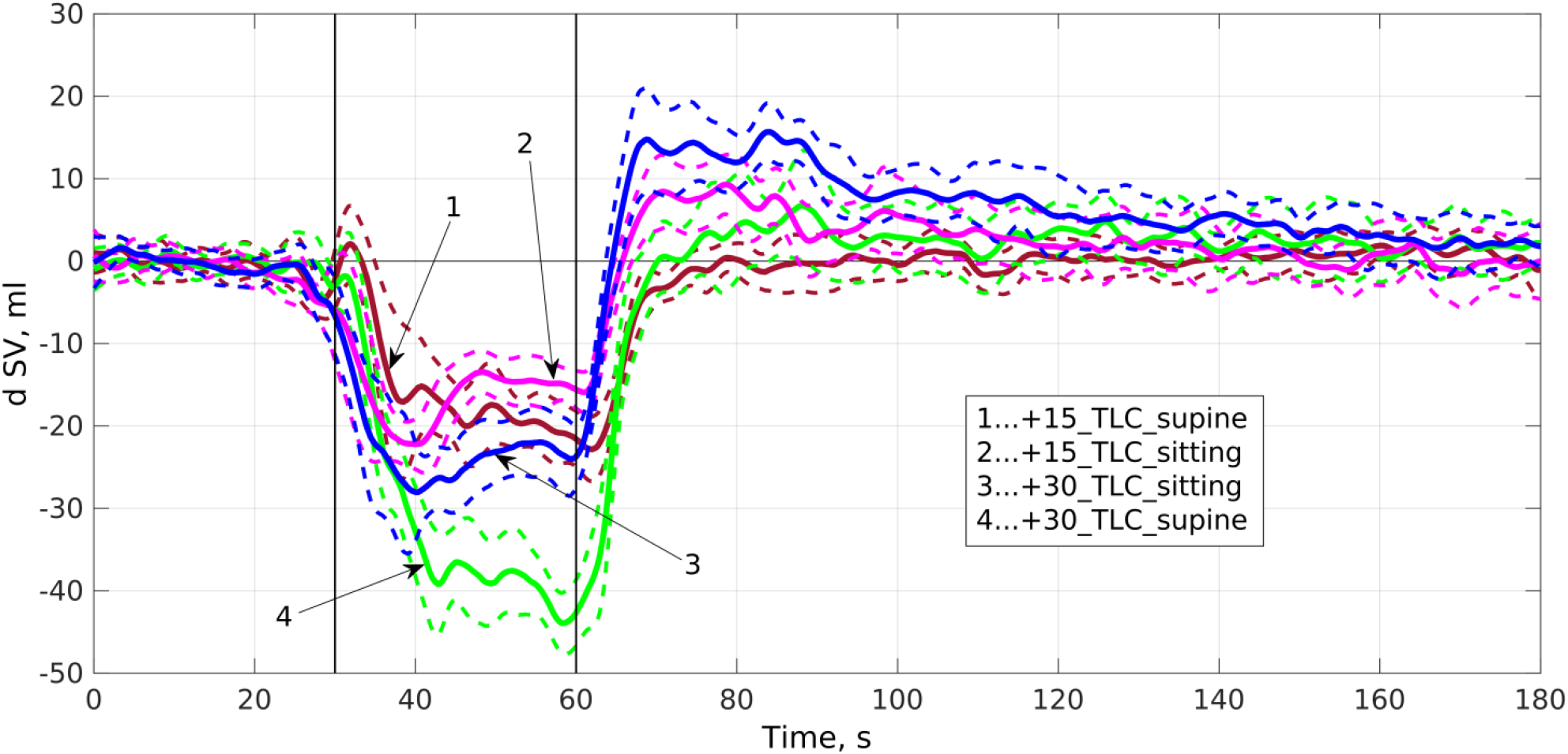
Example of simple parameter dynamics when performing a breathing maneuver. Stroke volume (SV) dynamics during +30_TLC and +15_TLC maneuvers is presented. Group I average curves are shown. Vertical lines indicate the start and the end points of the maneuver. Dashed lines correspond to the ±SD range. Prefix “d” denotes parameter changes from its baseline value.

For most of the considered physiological parameters, body position and lung volume did not change the direction of deviations from their baseline observed during maneuvers. With changing in lung volume or body position, deviation values did not change much. The main parameter determining the response of CVS during the maneuvers was alveolar pressure, the magnitude and sign of its change relative to atmospheric pressure. Figs. 3–8 show average (during the maneuver) deviations of physiological parameters from their baseline values.

**Fig. 3.**
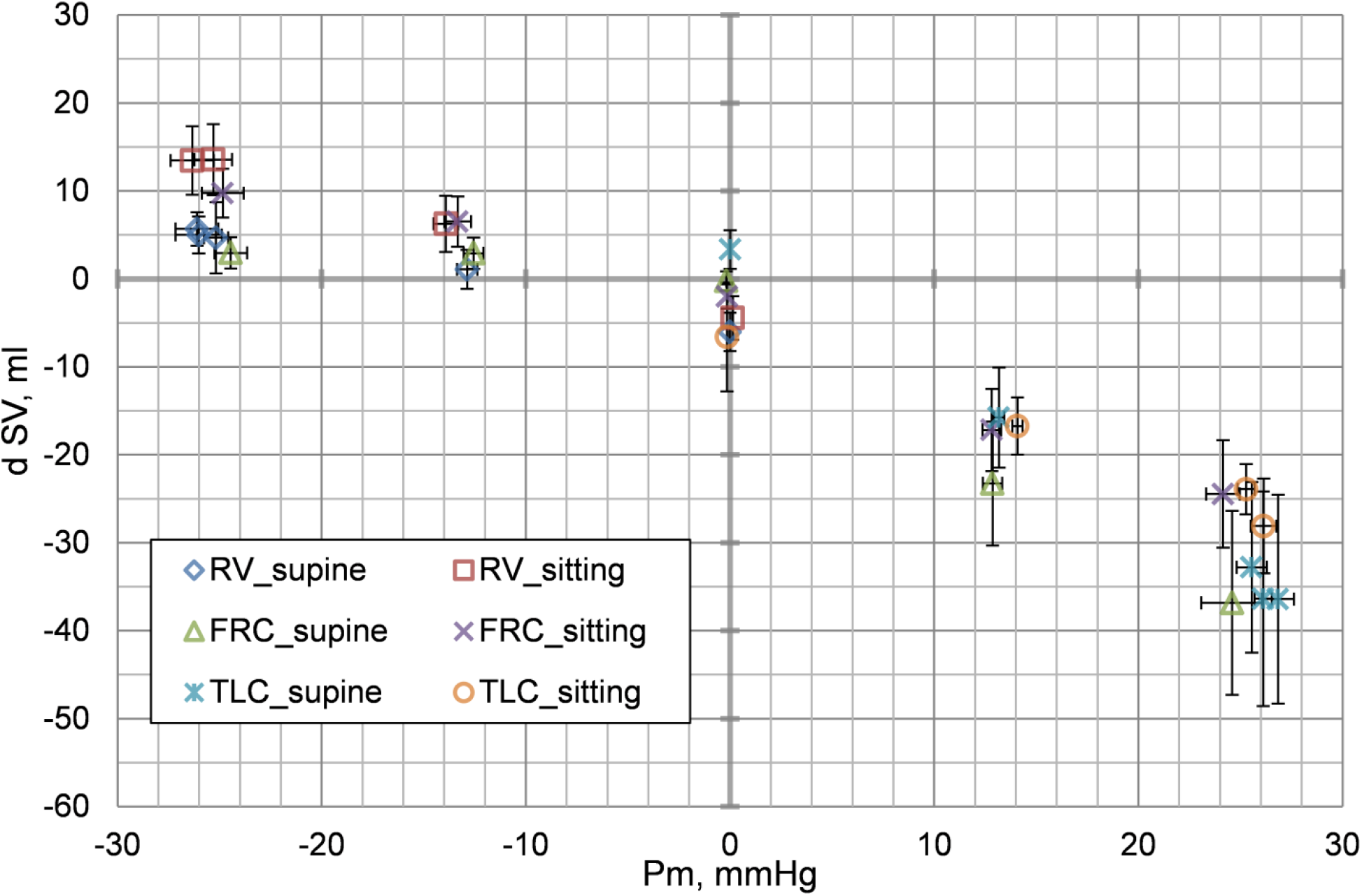
Mean changes in stroke volume (SV) during respiratory maneuvers. Pm – mouth pressure (relative to atmospheric pressure). Points for the same respiratory maneuver and close alveolar pressures refer to different volunteer groups (study series). Prefix “d” denotes parameter changes from its baseline value, whiskers represent standard deviation.

When performing maneuvers with any combination of lung volume, alveolar pressure and body position, periods of isovolumetric contraction and ventricular ejection changed inversely. As a result, their sum, equal to systole duration, practically did not change. The largest deviation did not exceed 10– 15% (about 0.03 s) of the baseline value.

Stroke volume (SV) decreased in maneuvers with positive alveolar pressure and slightly increased in maneuvers with negative pressure (Fig. 3), changes ranged from -35 to +15 ml. We should note that earlier in the study of the Müller maneuver effects on hemodynamics and the role of cardiopulmonary interactions, the opposite reaction was also observed: there was a decrease in SV and cardiac output (CO) during the maneuver [Condos et al., 1987]. The reason for the discrepancy in results can be the essentially different standardization of the maneuver: in the mentioned work, the decrease in airway pressure during the maneuver was at least 50 cmH_2_O for at least 10 s. In such conditions, rapid transients give a notable contribution to the observed result. In our study we tried to get information primarily about the mode that is established closer to the completion of the maneuver (when considering the Valsalva maneuver, late phase 2 – “pressure recovery”).

In maneuvers with negative alveolar pressure, HR practically did not change during maneuvers (Fig. 4). In maneuvers with positive pressures, HR increased up to +20 bpm.

**Fig. 4.**
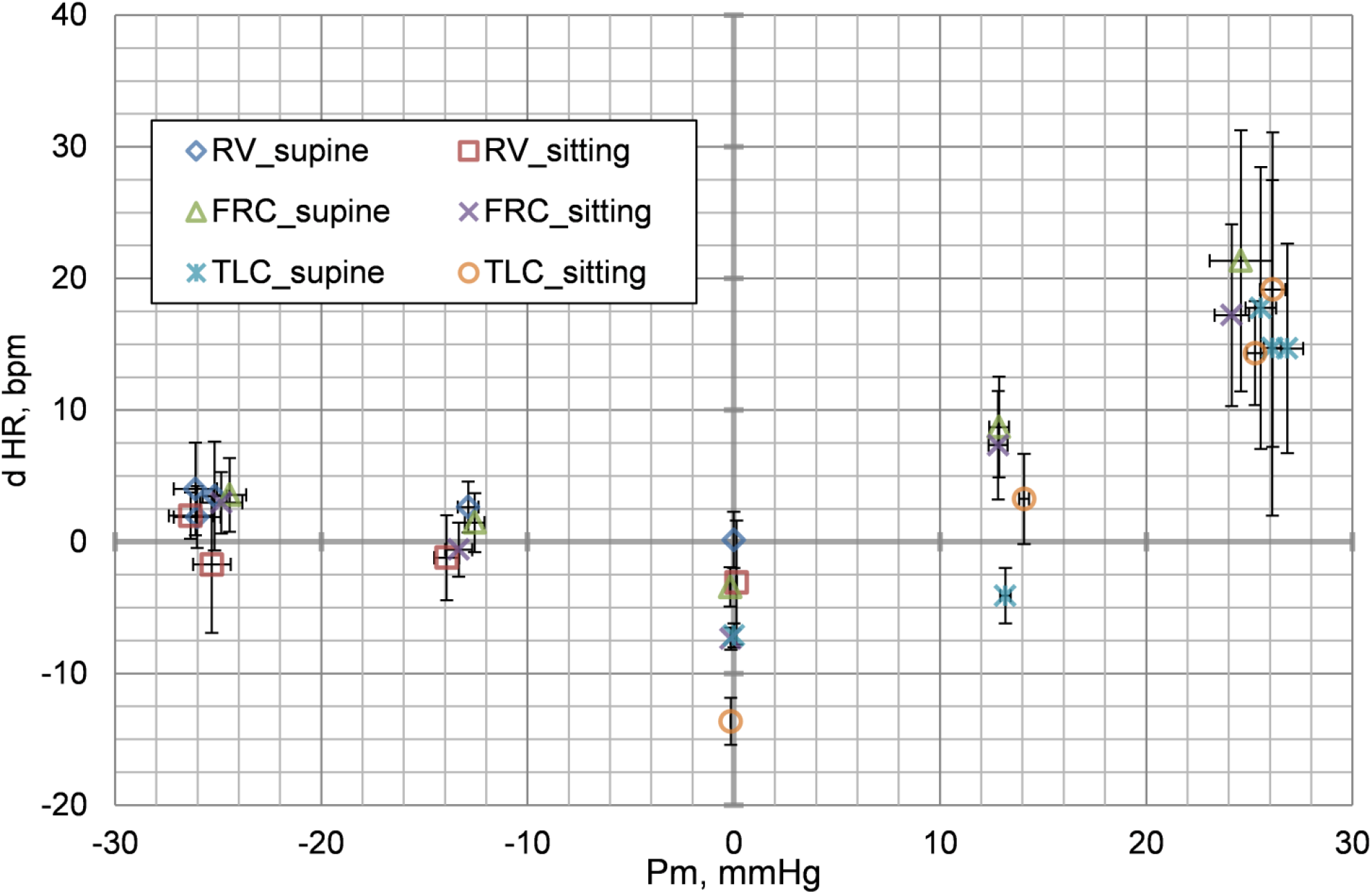
Mean changes in heart rate (HR) during respiratory maneuvers. Pm – mouth pressure (relative to atmospheric pressure). Points for the same respiratory maneuver and close alveolar pressures refer to different volunteer groups (study series). Prefix “d” denotes parameter changes from its baseline value, whiskers represent standard deviation.

Mean arterial pressure (MAP) decreased at decreasing mouth pressure (Pm) regardless of lung volume and body position; at increasing Pm, the result depended on lung volume (Fig. 5). When performing maneuvers at total lung capacity, MAP remained below baseline values, in other cases it increased. Changes in mean arterial pressure were within ±20 mmHg.

**Fig. 5.**
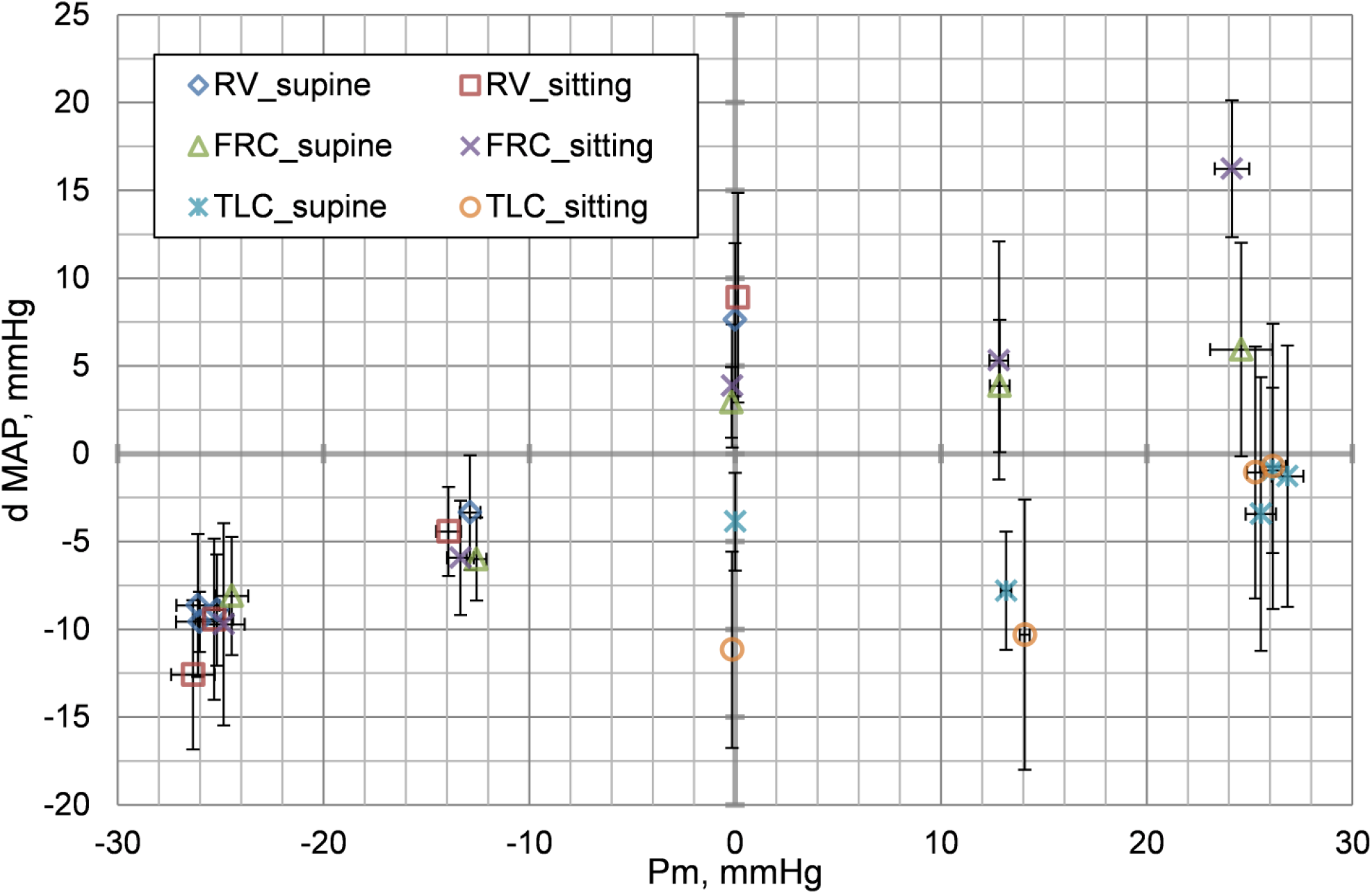
Mean arterial pressure (MAP) changes during respiratory maneuvers. Pm – mouth pressure (relative to atmospheric pressure). Points for the same respiratory maneuver and close alveolar pressures refer to different volunteer groups (study series). Prefix “d” denotes parameter changes from its baseline value, whiskers represent standard deviation.

Regardless of lung volume and body position, total peripheral resistance (TPR) decreased with decreasing Pm and increased with increasing Pm (Fig. 6). The range of changes in TPR was -0.3 to +0.7 mmHg·s/ml.

**Fig. 6.**
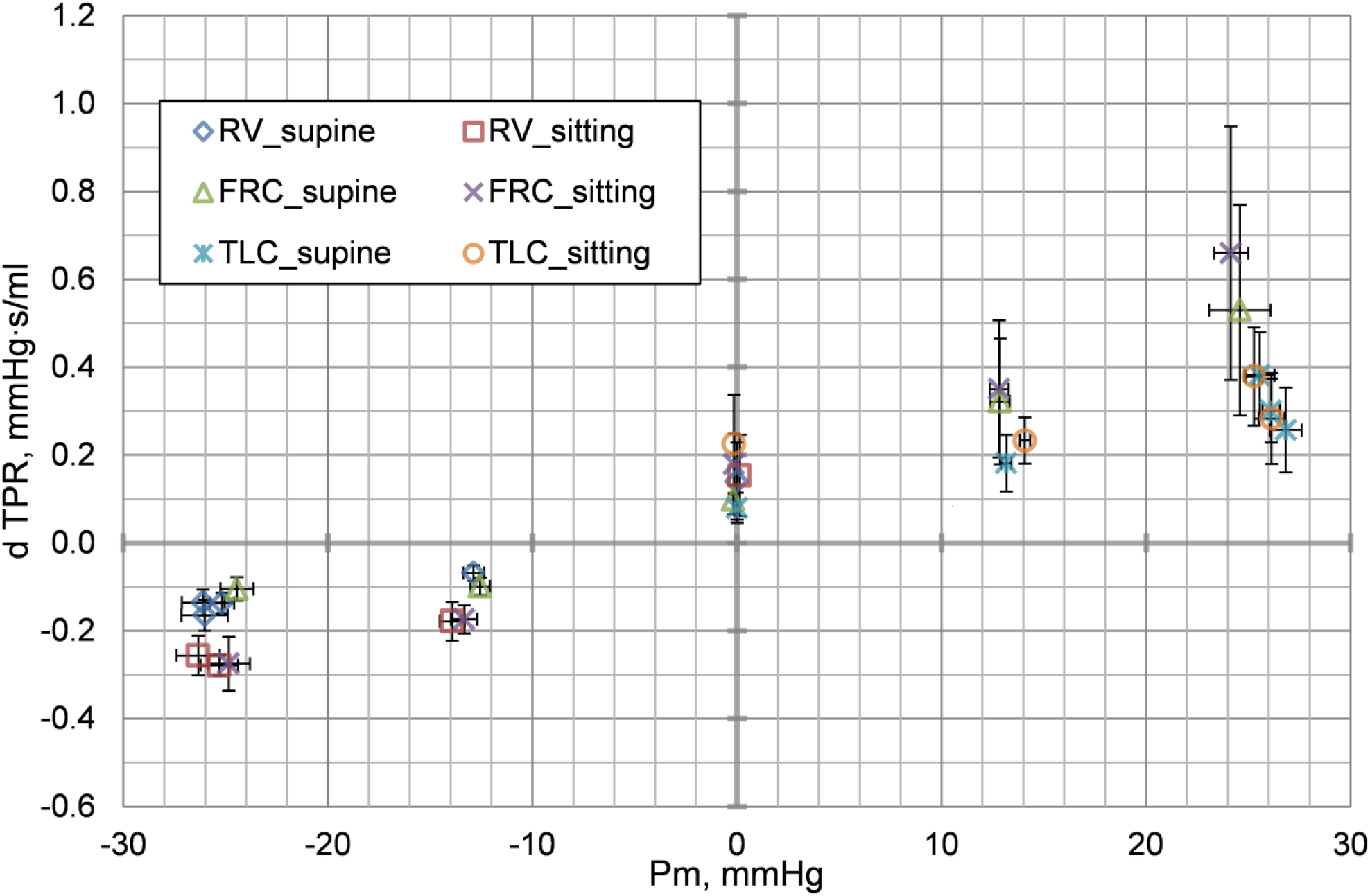
Mean changes in total peripheral resistance (TPR) during respiratory maneuvers. Pm – mouth pressure (relative to atmospheric pressure). Points for the same respiratory maneuver and close alveolar pressures refer to different volunteer groups (study series). Prefix “d” denotes parameter changes from its baseline value, whiskers represent standard deviation.

Cardiac output (CO) increased with decreasing alveolar pressure (Fig. 7). The magnitude of the increase was almost independent of body position and lungs volume, changes ranged from -2.0 to +1.25 l/min. As alveolar pressure increased, CO decreased despite the increase in HR. Changes in CO was determined by changes in SV.

**Fig. 7.**
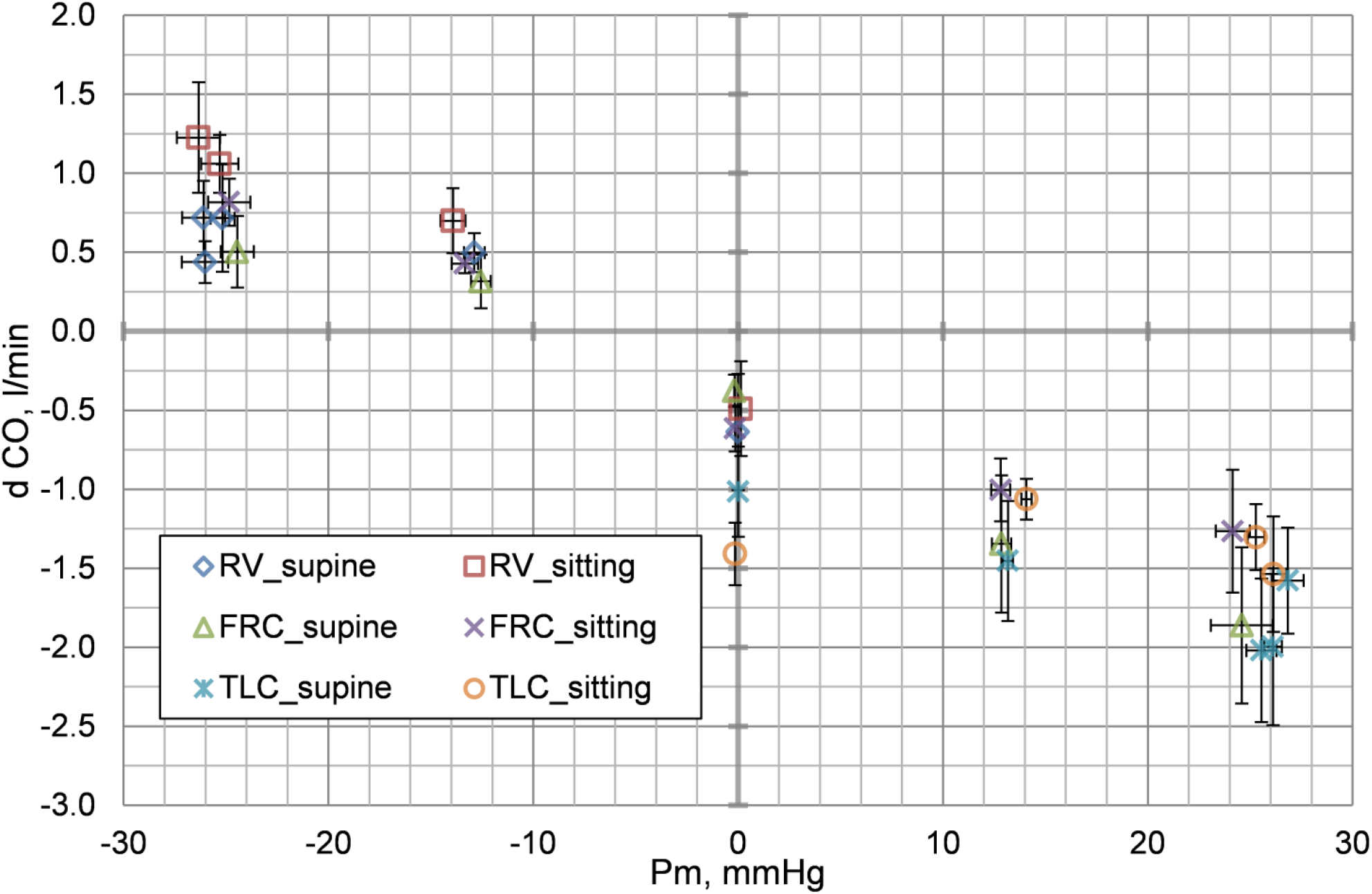
Mean changes in cardiac output (CO) during respiratory maneuvers. Pm – mouth pressure (relative to atmospheric pressure). Points for the same respiratory maneuver and close alveolar pressures refer to different volunteer groups (study series). Prefix “d” denotes parameter changes from its baseline value, whiskers represent standard deviation.

A decrease in baroreflex sensitivity (BRS) in the range of 2–8 ms/mmHg was detected during any maneuver. A more pronounced decrease in BRS (by 14 ms/mmHg) was observed only during the +30_TLC maneuver in a supine position.

Real parts of right thorax impedance measured at frequencies 5 and 50 kHz changed during maneuvers in the same way, curves almost coincided, the form of changes was a trapezoid. Imaginary parts of impedance practically did not change during maneuvers. Mean changes in R5/R50 during respiratory maneuvers are presented on Fig. 8.

**Fig. 8.**
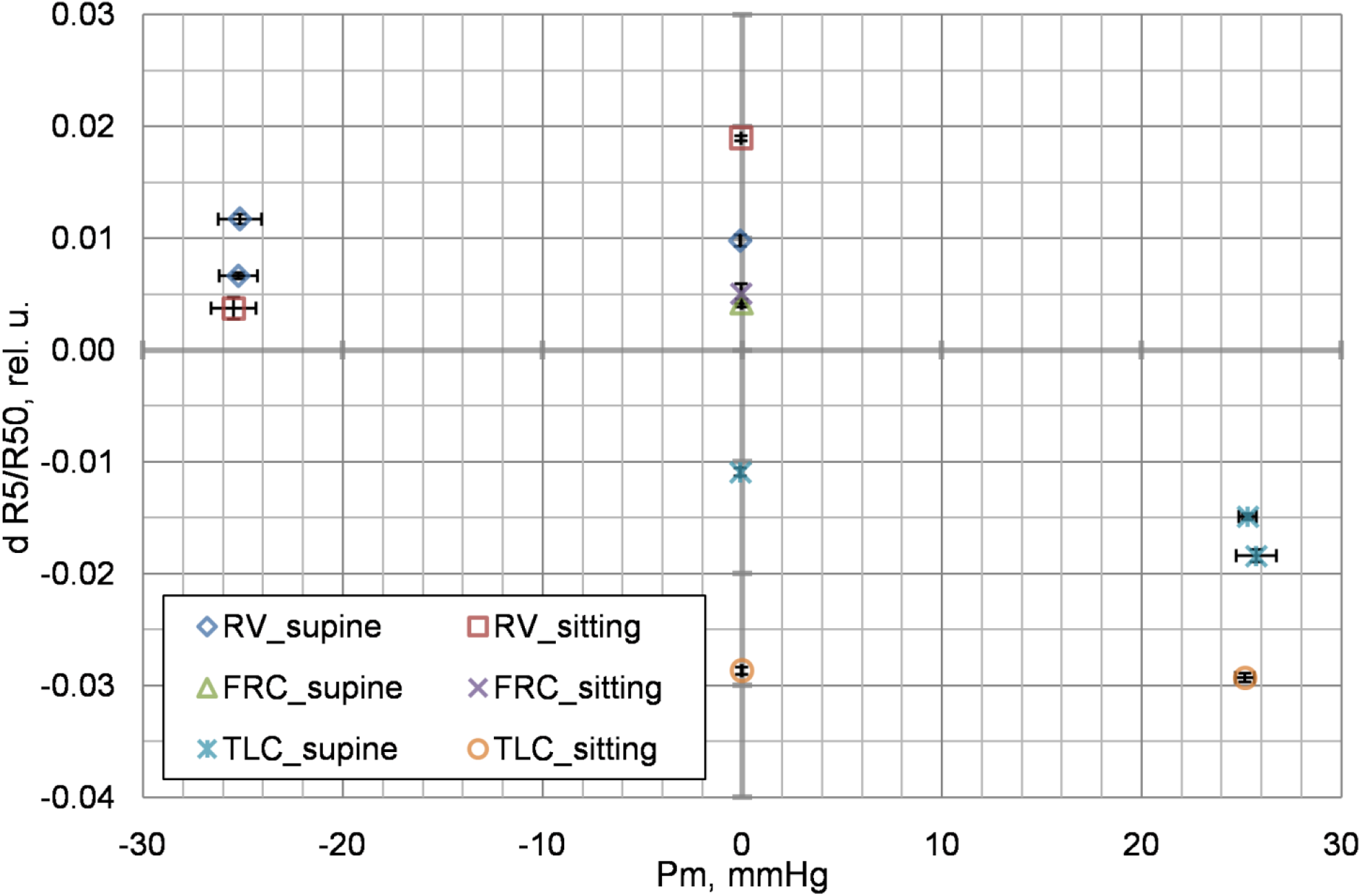
Mean changes in the ratio of real parts of right thorax impedance measured at 5 and 50 kHz (R5/R50, lung hydration index) during respiratory maneuvers. Pm – mouth pressure (relative to atmospheric pressure). Points for the same respiratory maneuver and close alveolar pressures refer to different volunteer groups (study series). Prefix “d” denotes parameter changes from its baseline value, whiskers represent standard deviation.

Changes in R5/R50 ranged from -0.03 to +0.02. Lungs volume caused the most pronounced effect on R5/R50. For maneuvers performed at RV or TLC in a sitting position, changes in R5/R50 were 2-3 times bigger than in a supine position. Positive alveolar pressure almost did not affect R5/R50 changes. In a supine position, negative alveolar pressure did not affect R5/R50 changes, but dramatically decreased changes in R5/R50 in a sitting position in maneuver performed at RV.

## Discussion

SV decreased in maneuvers with positive alveolar pressure and slightly increased in maneuvers with negative pressure (Fig. 3). Changes in SV indirectly indicate a change in cardiac end-diastolic volume when intrathoracic pressure changes. Stroke volume through the Frank-Starling mechanism is associated with end-diastolic volume. Filling of heart chambers during diastole occurs passively and depends upon differences of pressure inside the chambers and external (intrathoracic) pressure changing in accordance with alveolar pressure. Therefore, a decrease in alveolar pressure with unchanged duration of diastole and pressures in heart chambers results in an increase in end-diastolic and stroke volumes. An increase in alveolar pressure leads to their observed decrease.

In maneuvers with negative alveolar pressure, the increase in SV was less than its decrease in maneuvers with positive pressure. Asymmetry of SV changes may be caused by asymmetry of HR changes. In addition to intrathoracic pressure, two parameters can influence the value of SV changes: diastole duration and pressures in heart chambers during the diastole, which are close to pressures in large systemic and pulmonary veins. If the systole duration is practically unchanged, the diastole duration is determined by HR. In maneuvers with negative alveolar pressure, HR practically did not change during maneuvers (Fig. 4). Thus, there remain only two parameters affecting the magnitude of SV changes in these maneuvers: pressure in main veins and intrathoracic pressure. One can expect that vein pressures and intrathoracic pressure change symmetrically when Pm changes its sign. So in maneuvers with positive pressures, as HR increased, duration of diastole decreased, this additionally decreased end-diastolic volume and caused asymmetry of SV changes.

One would expect that for maneuvers with negative alveolar pressure, a decrease in pressures in main veins would be approximately the same in magnitude as a decrease in intrathoracic pressure. In this case, the pressure difference stretching heart chambers during diastole would not change, end-diastolic and stroke volumes would not change either. However, the increase in SV observed in the study is only possible with an increase in transmural pressure acting on ventricles during diastole. Hence, a decrease in intrathoracic pressure is greater than a decrease in pressure in pulmonary veins and intrathoracic sections of main veins of systemic circulation. Besides, when alveolar and intrathoracic pressures decrease, one can expect a decrease in the resistance of pulmonary vessels associated with their stretching. Lowering resistance of pulmonary vessels also contributes to SV increase.

Greater changes in SV were observed for any of lung volumes for maneuvers performed in a sitting position than in a supine position. Besides, in a supine position, changes in SV did not depend on the decrease in alveolar pressure at any lung volume, while in a sitting position, a two-fold increase in alveolar pressure drop resulted in an increase in SV by approximately 1.5–2 times regardless of the lung volume. This peculiarity can be explained from the point of view of gravitation-dependent non-uniformity of blood flow in lungs. In a supine position, blood flow in any part of lungs is approximately equal, blood vessels are stretched. In a sitting position, due to hydrostatic pressure gradient, negative transmural pressure affects some vessels. These vessels are in a partially or completely collapsed state, blood flow through them is reduced or absent at all. In a supine position, a decrease in alveolar pressure may lead only to a slight increase in the lumen of pulmonary vessels and a corresponding decrease in their resistance. In a sitting position, the reduction of alveolar pressure increases transmural pressure acting on lung vessels, partially compensating for uneven distribution of blood flow, some fully collapsed vessels will be stretched and included in blood circulation, which will notably reduce total resistance of pulmonary vessels. The proportion of vessels involved in blood circulation will increase approximately in proportion to the value of a decrease in alveolar pressure until all pulmonary vessels are straightened. It is possible to estimate the magnitude of the necessary reduction in pressure. Since the distance from heart level to the tops of the lungs is about 20 cm, to compensate for blood flow unevenness, it is necessary to create a pressure reduction of about 20 cmH_2_O. In a sitting position and under normal gravity, full compensation of the unevenness can be expected when alveolar pressure is reduced by 20–40 cmH_2_O below the atmospheric pressure, as about half of this pressure reduction will be compensated by tissues stretching [Tikhonov et al., 2003]. Experimental data confirm this estimation [Bake, 1971].

The decrease in TPR associated with a decrease in MAP (Fig. 5, Fig. 6) and practically unchanged HR (Fig. 4) in maneuvers with negative alveolar pressure might be a consequence of oppositely directed reflex influences. When alveolar pressure decreases, MAP decreases, so carotid receptors stimulation decreases, while it possibly increases for aortic arch receptors. For mentioned reflexogenic zones, an increase in receptors stimulation leads to bradycardia and a decrease in TPR. With a decrease in stimulation, the opposite effects are observed. For aortic baroreceptors, one may expect an increase in stimulation, as the decrease in MAP is less than the decrease in intrathoracic pressure even taking into account the approximately two-fold attenuation [Tikhonov et al., 2003] in relation to changes in alveolar pressure. It follows from the data [Smith et al., 2001] that in humans aortic baroreflex prevails over carotid baroreflex. So observed with negative alveolar pressure paradoxical decrease in TPR, while MAP is also decreasing, may be explained by assuming that transmural pressure for aortic arch increases when intrathoracic pressure decreases. Also lack of HR response to decrease in alveolar pressure, while the response to positive alveolar pressure is pronounced, may be speculatively explained by increasing aortic arch transmural pressure and oppositely directed reflex influences.

An increase in CO_2_ tension in the blood during respiratory maneuvers may also be the cause of changes in TPR. However, almost complete absence of changes in HR in maneuvers with negative alveolar pressure and the lack of dependence of HR and TPR changes on the lung volume, as well as the reverse of changes in TPR when changing the sign of alveolar pressure, indicates a minor contribution of this mechanism of vascular tone regulation to the observed responses during 30-second apnea.

The vascular response to change in shear stress at the vessel wall caused by change in the magnitude of blood flow could influence TPR. Within the results of this study, it is impossible to separate the influence of baroreflexes and local vascular response to shear stress changes.

Changes in MAP during maneuvers are result of blood flow mechanics and action of plenty regulatory mechanisms controlling circulation. Within the results of the study, it is impossible to separate their contribution to observed changes in MAP as well as to changes in HR.

Changes in R5/R50 at changes in lung volume and zero (i.e. equals to atmospheric) alveolar pressure (Fig. 8) are easy to explain: changing lung volume leads not only to changes in the amount of gas in lungs but also to changes in the amount of blood. At deep expiration, lung vessels are “squeezed” by increased external (intrathoracic) pressure; at deep inspiration, the opposite process takes place. Blood volume changes affect R5/R50 value more than gas volume changes. In a sitting position, the magnitude of changes is greater due to a different position of diaphragm and different intrathoracic pressure. The points reflecting 0_FRC maneuvers do not equal exactly to zero, because free breathing is chosen as baseline. Average lung volume during free breathing is slightly higher than FRC.

In a supine position, when alveolar pressure decreases, R5/R50 value changes in the same way as in the maneuver with alveolar pressure equal to atmospheric pressure. It indicates that in a supine position lung vessels are already stretched, they cannot change the volume when transmural pressure further increases. For similar maneuvers performed in a sitting position, there is a decrease in R5/R50, indicating an increase in blood filling with an increase in transmural pressure (partially collapsed vessels are distending).

In a supine position, when alveolar pressure increases in maneuver at TLC, some additional decrease in R5/R50 is observed, lung blood filling increases. This phenomenon can be explained by changes in intra-abdominal pressure during maneuvers performing. In maneuvers with positive alveolar pressure, abdominal muscles actively participate and diaphragm is passive (the situation is reversed in RV-maneuvers). Involving abdominal muscles lead to increase in intra-abdominal pressure together with intrathoracic pressure. Taking into account that a large part of blood volume is concentrated in abdominal veins, the increase in intra-abdominal pressure will lead to “extruding” of blood into other body parts, including thorax. It seems that in maneuvers with positive alveolar pressure, the increase in intra-abdominal pressure is slightly greater than the increase in intrathoracic pressure, so some amount of blood is redistributed from abdominal vessels to the thorax.

Thus, the observed changes in all measured parameters allow us to identify two most likely mechanisms determining the response of CVS to a short-term decrease in intrathoracic pressure. The first is pulmonary vessels stretching induced by intrathoracic pressure reduction, which reduces resistance of the vessels, increases their blood filling, and partially compensates for hydrostatic pressure difference in the vessels between different parts of lungs. This effect seems to affect mainly small lung vessels providing over 90% of total resistance of pulmonary circulation [Dyachenko and Shabelnikov, 1985]. The second is the increase in SV by the Frank-Starling mechanism. In case of a short-term increase in intrathoracic pressure, main operating mechanisms determining CVS response, apparently, remain the same, but the direction of changes in CVS parameters is opposite.

When considering the response of CVS to intrathoracic pressure variations, special attention should be paid to the fact that, in some cases, aortic and carotid baroreflexes may have an opposite effect on vascular tone.

Besides, when analyzing the response of CVS to respiratory maneuvers with increased airway pressure, one should consider the increase in intra-abdominal pressure, which can notably change the distribution of blood between abdominal and thoracic regions. In maneuvers associated with a decrease in airway pressure, intra-abdominal pressure changes little due to mechanics of breathing. In this case, the effect of intra-abdominal pressure on CVS may not be considered.

## Conclusion

We obtained quantitative information about the response of human circulation to changes in alveolar pressure at different lung volumes and to changes in lung volumes at unchanged alveolar pressure in supine horizontal and sitting positions. Changes in alveolar and intrathoracic pressures influenced hemodynamic parameters stronger than changes in lung volume or body position. SV increased when alveolar pressure decreased and decreased when pressure increased irrespective of lung volume and body position. The changes ranged from -35 to +15 ml. In a sitting position, the effect of alveolar pressure changes was more pronounced. HR virtually did not change with decreasing alveolar pressure, but increased with increasing pressure (up to +20 bpm). MAP decreased at decreasing alveolar pressure regardless of lung volume and body position; at increasing pressure, the result depended on lung volume. When performing maneuvers at TLC, MAP remained below baseline values; in other cases, it increased. MAP changes were within ±20 mmHg. Regardless of lung volume and body position, TPR decreased with decreasing alveolar pressure and increased with increasing alveolar pressure; the range of changes was from -0.3 to +0.7 mmHg·s/ml.

## Acknowledgments

We sincerely thank for their help and friendly support our colleagues from the State Research Center of the Russian Federation – Institute of Biomedical Problems of the Russian Academy of Sciences: PhD in Medicine Dr. Galina Reushkina, PhD in Medicine Dr. Julia Popova, PhD in Medicine Dr. Dilyara Khusnutdinova, and Alexander Zolotarev. We would also like to acknowledge the invaluable contribution of the late PhD in Medicine Dr. Alexander Suvorov.

The work was partially supported by the Russian Academy of Sciences, Program “Integration of Control in ensuring the functions of the body” (grant IV.7.1.) and by the Program of basic research of the State Scientific Center of the Russian Federation – Institute of Biomedical Problems of the Russian Academy of Sciences (grant FMFR-2024-0038).

## Declaration of Competing Interest

The authors declare that they have no known competing financial interests or personal relationships that could have appeared to influence the work reported in this paper.

## Notes

### Competing Interest Statement

The authors have declared no competing interest.

